# The Barcode Inference Pipeline (BIP): From Sequencer Output to DNA Barcodes

**DOI:** 10.64898/2026.07.21.739876

**Authors:** Sean WJ Prosser, Ken A Thompson, Nicholas W Bard, Robin M Floyd, Emine Ozsahin, Paul DN Hebert

## Abstract

DNA barcoding involves the recovery of a DNA sequence for a target gene region from its source specimen. This process becomes complex when multiple sequences are recovered from a specimen. This diversity can reflect methodological artifacts (e.g., chimeras, PCR errors, sequencing errors, tag jumps) and real template diversity in the DNA extract (e.g., contamination, endosymbionts, nuclear-mitochondrial pseudogenes (NUMTs), parasites). To support the analysis of sequence data from millions of specimens annually, the Centre for Biodiversity Genomics (CBG) has developed BIP, the Barcode Inference Pipeline. Compatible with all major sequencing platforms, BIP processes fastq files and returns both target DNA barcodes and non-target sequences (NTS). BIP implements filtration, demultiplexing, primer trimming, read denoising, chimera scanning, sequence error correction, operational taxonomic unit (OTU) delineation, OTU taxonomy assignment, and selection of the target barcode. When analysis examines the cytochrome *c* oxidase 1 (COI) barcode region, BIP assigns each OTU to a Barcode Index Number (BIN) when there is a match. BIP also returns summary files, including a taxonomic assignment for each OTU based on a DNA barcode reference library. We describe BIP’s flexibility and structure before demonstrating its functionality by analyzing COI sequence data generated from both nanopore and Illumina platforms, as well as rbcLa and ITS2 data generated from plants, and 18S data generated from arthropods. Because it disambiguates target and non-target sequences, BIP outperforms an alternative pipeline package, ONTbarcoder. To ease access, installation, and functionality, BIP is provided as a Docker container and is also available in the Galaxy toolshed.

## INTRODUCTION

Over the past decade, DNA sequencing has transitioned from the most to the least expensive component of DNA barcode analysis (Hebert *et al*. 2025). Today, researchers routinely sequence specimens in their own laboratories (Sánchez-Vendizú *et al*. 2025). The ability to generate sequence data (Srivathsan *et al*. 2024) has created the need for accessible, reproducible, and easily implemented bioinformatic workflows.

Bioinformatic analysis for DNA barcoding—the practice of generating short sequences from a specific region of the genome for species identification (Hebert *et al*. 2003)—follows a standard workflow. Analysis begins by removing reads with low quality scores, those outside the expected size range, and those where the unique molecular identifiers (UMIs) and primers used to tag amplicons from different specimens cannot be unambiguously discriminated (Martin 2011, Wick, Judd, and Holt 2018). Next, reads are clustered into operational taxonomic units (OTUs) employing user-defined thresholds (Callahan *et al*. 2016). Finally, these OTUs gain a taxonomic assignment based on their comparison with a reference library for the target gene region (Porter and Hajibabaei 2022, Li *et al*. 2024, Furneaux *et al*. 2025). Because its components are standardized, this workflow is amenable to deployment within user-friendly distributed pipelines. However, many current pipelines lack critical steps such as demultiplexing or taxonomic assignment, opting to leave these tasks to the user as separate pre- or post-analysis requirements (Hakimzadeh *et al*. 2024). Furthermore, most existing pipelines were built for metabarcoding—where multiple OTUs are in each sample—which introduces overlooked complexities when applied to single-specimen analysis. The barcode workflow often yields multiple OTUs per specimen (Hebert *et al*. 2025) due to co-amplification of biologically associated organisms (e.g., endosymbionts and parasites), exogenous contaminants, or artifactual sequences (e.g., chimeras), and the user must determine which, if any, is the target barcode sequence. While existing pipelines can identify all OTUs associated with a single specimen, none infers which OTU is most likely to be the target barcode despite having access to the necessary information: specimen taxonomy, OTU sequence length, OTU sequence quality, and OTU read depth. Indeed, a “full-service” bioinformatic pipeline for DNA barcoding—one that ingests raw sequence data in a platform-agnostic, marker-agnostic, and primer-agnostic manner and designates all OTUs as either target or non-target sequence (NTS)—does not exist.

To address this gap, we release the bioinformatic pipeline used at the Centre for Biodiversity Genomics (CBG)—termed the Barcode Inference Pipeline (BIP)—which underpins production at the world’s largest DNA barcoding facility. BIP is distinguished from alternative pipelines in several ways. It ingests raw sequence data and returns high-quality DNA barcode sequences, without pre- or post-analysis manipulation. BIP also accepts data from second- and third-generation sequencing platforms—short Illumina paired-end reads (Meier *et al*. 2016), long PacBio circular consensus sequences (Hebert *et al*. 2018), or long nanopore reads (Hebert *et al*. 2025). It can, as well, analyze any molecular marker including those generated by multi-marker or multi-primer studies. BIP returns both target and non-target sequences (NTS), providing the capacity to probe the origins of this diversity (i.e., random contamination versus species interactions) (Baker *et al*. 2016). BIP contains features designed to optimize the fidelity of results when analysis targets the 5’ end of the cytochrome *c* oxidase subunit I (hereafter COI) marker, the standard barcode for the animal kingdom and the most widely used marker on the Barcode of Life Data System (BOLD). These COI-specific features include polymerase-error correction and Barcode Index Number (BIN)-matching (Ratnasingham and Hebert 2013), allowing users to match their results to known species proxies. The underlying logic behind BIP has been used to recover barcodes from millions of specimens at the CBG, so its performance has been evaluated across many taxonomic groups. Nonetheless, this publication demonstrates BIP’s functionality by analyzing COI, rbcLa, ITS2, and 18S sequence data generated from either nanopore or Illumina platforms.

## METHODS

### Installation and use

BIP is installed using the image file in the Docker Hub Container Image Library (https://hub.docker.com/). Running a container from the image file generates BIP’s working environment and installs the appropriate versions of all software required for its operation. First-time users can execute a standardized command to analyze the trial dataset in the container to confirm successful installation as described in the documentation. Detailed operating instructions are available at github.com/cbg-innov/BIP. To initiate a run, BIP requires at least one gzipped .fastq file (or two files—R1 and R2—if using Illumina paired-end data), an Excel file (.xlsx) with parameters, and a reference library (.fasta). BIP installs quickly on Linux, macOS, and Windows. We note that all technical descriptions in this manuscript reflect BIP v1.1.0 and the most current descriptions will always be on GitHub.

In the BIP parameters file, one tab allows the user to enter sample-specific information: a unique specimen identification code, plate and well locations (used to generate visual heat maps of the results), the nucleotide sequences of the unique molecular identifiers (UMIs) used to index each sample, gene-specific primer names, whether a sample is a negative control, and the taxonomic identification—kingdom down to species, where known—of the samples. Another tab allows the user to enter primer-specific information: primer nucleotide sequences, locus name, reference library file name, minimum, maximum, and mean amplicon length, denoising and OTU clustering parameters, and taxonomic assignment thresholds. These data allow BIP to distinguish target from non-target sequences. For instance, if the user specifies that a specimen belongs to the order Hemiptera and valid sequences are recovered from two orders—Diptera and Hemiptera—the hemipteran sequence is designated as the target while the dipteran sequence is categorized as a non-target sequence (NTS).

BIP supports taxonomic reference libraries in sintax-formatted fasta files (Rognes *et al*. 2016). In sintax format, the sample name is followed by specifically structured taxonomic information. Optionally, BIN designations can be included following a vertical bar (i.e., ‘|’) after the sample ID—required for BIN matching in COI runs. To meet this need, BOLDistilled reference libraries (Prosser *et al*. 2025) are offered in this format. An example of a valid header is:

>Example_SampleID|BOLD:AAA0001;tax=k:Animalia,p:Chordata,c:Mammalia,o:Primates,f:Hominidae,g: Homo,s:Homo sapiens

Any custom local reference library provided in sintax format can be referenced by file path in BIP—it does not require additional installation. Any number of additional target genes and reference libraries can be used in a run and need only be indicated by the user in the input parameters file.

### Workflow

BIP’s workflow is illustrated in Fig. 1. BIP is pointed towards a working directory containing the input data. For long-read sequencers (e.g., PacBio, Oxford Nanopore), BIP accepts one or more gzipped .fastq files as input. Paired-end (i.e., Illumina) data requires one gzipped R1 and one gzipped R2 file. The user must specify their data type when using BIP. Most base-calling algorithms include basic read filtering based on mean quality scores, which are calculated in a platform-specific manner. BIP therefore does not re-filter reads on quality alone. Instead, it excludes low-quality reads indirectly by first removing reads outside the user-specified length range and then only retaining reads if both gene-specific primers are found. These criteria exclude reads with high levels of substitution, insertion, or deletion errors. Reads that survive both length filtration and the check for primer integrity are demultiplexed with *cutadapt* (Martin 2011) by examining their UMIs. If symmetrical UMIs are detected (i.e., the same UMI sequence was used to tag each end of the amplicon), BIP will identify and remove tag-switched reads (Schnell, Bohmann, and Gilbert 2015). Next, primers and UMIs are trimmed, leaving only the target gene region.

**Figure 1:**
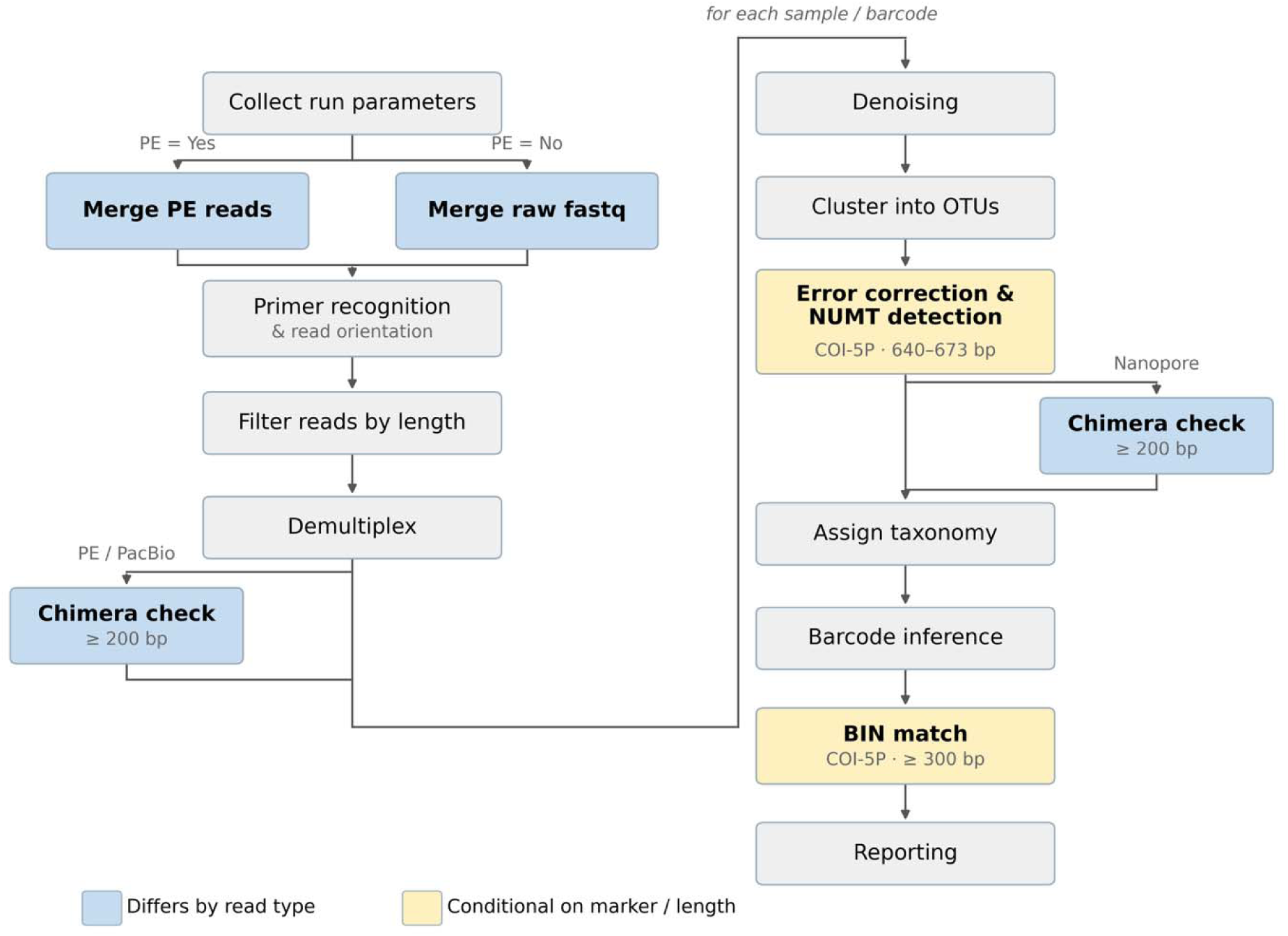
Diagram of the BIP workflow. PE = paired-end reads.

The next step is either chimera removal (vsearch --uchime_denovo) using *vsearch*’s *de novo* chimera detection (Rognes *et al*. 2016) or OTU clustering (vsearch -- cluster_fast), depending on the type of data being analyzed. Chimeras form during PCR when the strands of two genetically distinct amplicons anneal sufficiently for a polymerase to bind and elongate, resulting in a chimeric amplicon composed of parts of each source (Meyerhans, Vartanian, and Wain-Hobson 1990). Chimeras can usually be recognized by their low abundance relative to their source molecules (Edgar *et al*. 2011). *De novo* chimera detection relies on comparing low-abundance molecules to high-abundance molecules from the same source well to see if the former could have arisen from a combination of the latter. It is therefore best to perform this screen early in the bioinformatic pipeline before too many reads are discarded; however, sequencing errors will confound *de novo* chimera detection and should be corrected first. To that end, for data with few or no sequencing errors (e.g., Illumina PE or PacBio), chimera removal occurs immediately after demultiplexing, as soon as reads are grouped with others from their own PCR. For error-prone nanopore data, chimera removal occurs later in the pipeline (see below).

Errors can occur during both PCR and sequencing. Errors during PCR can be present in both source and chimeric amplicons and are therefore tolerated by *de novo* chimera detection algorithms. By contrast, errors accrued during sequencing (i.e., after chimera formation) break the link between chimeras and their source molecules and must be corrected prior to *de novo* chimera detection. Since sequencing errors are mostly random noise (Teeling and Glöckner 2012), they can be mitigated by a process called denoising, often by clustering similar reads and generating a consensus sequence to mask the random errors. Demultiplexed reads are denoised in BIP using vsearch -- cluster_fast (Rognes *et al*. 2016) guided by a user-defined threshold. As this threshold depends on the marker being used, we suggest it be the maximum intra-specific distance of the marker gene plus the sum of the expected PCR error rate and sequencing error rate (e.g., ∼3% for COI plus 2% for PCR with a non-proofreading polymerase and nanopore sequencing, for a total of 5%). As a further quality control measure, OTUs are only retained if they reach a minimum read depth set by the user (we recommend starting with ≥ 3 reads regardless of the platform and adjusting thereafter). A second clustering step with a user-defined threshold (typically a higher percent identity closer to the intra-specific distance of the marker) is then employed to refine the OTUs. Consensus sequences are generated using a default 75% majority (this can be adjusted by the user), and ambiguous bases (Ns) are inserted where a consensus could not be reached.

If the primers generate full-length amplicons of COI, a custom error correction script is then applied (*homopolymer_error_fix.py*). It aligns the sequence of each OTU to a reference database that includes a large array of known amino acid sequences for COI; for more details on error correction, see the methods in the manuscript describing BIP’s metabarcoding companion software, MAP (Prosser *et al*. 2026).

Although random sequencing errors have been corrected by this point, polymerase errors generated during PCR, especially those generated during early cycles of PCR (Shagin *et al*. 2017), may persist. Since polymerase errors occur primarily in homopolymer tracts, and nanopore sequencing is prone to miscounting repeats of the same base within homopolymer tracts (Liu-Wei *et al*. 2024), the error correction algorithm targets these regions and identifies any 1–2 bp frameshift insertions/deletions (indels). It then adds ambiguous ‘N’ characters or deletes the excess base(s) to restore the reading frame. To avoid “correcting” true biological diversity, edits are only made if the indel: 1) lies within a homopolymer tract (i.e., known hot spots for such errors); and 2) is 1–2 bp in length (i.e., not natural codon indels). COI sequences are then checked for stop codons and residual frameshift indels; if a sequence is in-frame but contains stop codons, it is designated as a potential nuclear-mitochondrial pseudogene (NUMT). NUMTs are ancestral mitochondrial gene sequences which have been integrated into the nuclear genome (Lopez *et al*. 1994). Because they no longer function, they are free to acquire indels and/or stop codons. In-frame sequences with stop codons are tagged as likely NUMTs, while sequences with uncorrectable indels may reflect NUMTs, early-cycle PCR errors, or highly erroneous sequences; both categories are disqualified from being the target barcode.

Following error correction and NUMT-flagging, chimera removal proceeds here for nanopore data, and taxonomy is assigned probabilistically using *sintax* (vsearch -- sintax) (Rognes *et al*. 2016) with a user-defined confidence threshold. Briefly, assignment involves placing query sequence k-mers into a tree composed of reference sequence k-mers with known taxonomic hierarchy. Iterative placements track the location of each sequence in the guide tree and consolidate the results into probabilistic assignments at each rank. If the marker is COI (≥ 300 bp) and BINs are included in the reference library, BIP conducts BIN-matching using vsearch --usearch_global (Rognes *et al*. 2016). The percent identity value to return a BIN match is set by the user independently for each primer set.

OTUs are then designated as either ‘target’ or ‘non-target sequence’ (NTS) through a process we term ‘barcode inference’. A maximum of one target sequence (i.e., barcode) can be returned for a specimen, but there is no maximum number of NTS. An OTU must match the specimen’s assigned taxonomy and contain no known sequence issues (i.e., indels or stop codons for COI) to be designated as target. The user can set the taxonomic rank to enforce matching at a specified rank. For example, if a user is confident in their identifications to family, they can implement the flag, `-- min_rank_match f`, and then a mismatch at the family level will prevent assignment as target. Ranks use the same standard letters as *vsearch*’s sintax. Missing values are ignored, so if a specimen is missing a rank at the family level, it will be enforced at the order level if an identification is supplied at that rank.

If multiple OTUs meet the above ‘target’ criteria—or if no taxonomic information is provided by the user—tie-breaking is employed. As a first pass, BIP examines lower taxonomic ranks than the minimum, if they are provided by the user, to evaluate matching. For example, if `--min_rank_match` was set to order, but the user provided specimen data down to species, and one of the tied OTUs matches to the expected family/genus/species while the other only matches to order, the OTU matching at lower ranks is designated as the target barcode. If a target still does not emerge, the OTU closest to the expected sequence length is assigned as target. For example, if the expected sequence length is 658 bp, then a sequence of length 655 bp would be selected over a sequence of length 664 bp. If still tied, the OTU with fewer ambiguous bases (i.e., higher quality) is selected as the target sequence. If a tie still remains, the OTU with the higher read count—more likely to be the target (Hebert *et al*. 2025)—is designated as the target. In the unlikely event that two or more OTUs are still tied, all are designated as NTS.

Results are returned—separately for each marker in multi-marker runs—in formats that facilitate run assessment, publication/archiving, and flexible downstream analysis. A single summary report (.pdf) is generated that provides plots and metrics useful for assessing technical and biological aspects of the run (e.g., step-specific read counts, negative control summaries, non-target sequence metrics, barcode recovery rates, and various OTU metrics). Post-demultiplexed reads (.fastq) are provided for publication/archiving purposes. OTU component reads (.fasta) are provided so the user can assess the internal composition of an OTU. A detailed OTU table (.tsv; Supplemental Files S1–S4) is provided for human inspection or further analysis. It comprises one row per OTU with columns containing data such as the OTU consensus sequence, sequence length, ambiguous base count, read depth, “barcode status” (i.e., target or NTS), taxonomic assignments (from kingdom to species, where available) as well as the sintax-based probability scores, user-supplied taxonomy, and whether each rank matches the OTU’s assigned identity. The target barcode sequences, as well as NTS, are also provided (.fasta), but caution is advised when analyzing these sequences (see Supplemental Text for recommendations on how to interpret NTS data).

### Validation

In this publication, we demonstrate the performance and flexibility of BIP by presenting results generated by BIP v1.1.0 on sequence data from four runs involving different amplicon lengths, markers, and sequencing platforms (Table 1):

1. A large-scale (100,320 arthropod specimens), full-length (658 bp) COI-barcoding run sequenced on a MinION flow cell with a MinION Mk1B nanopore sequencer (ONT).
2. A medium-scale (380 arthropod specimens), short-read (418 bp) COI-barcoding run sequenced on a P1 flow cell with an Illumina NextSeq 2000 sequencer.
3. A small-scale (95 flowering plant specimens), full-length multi-marker (rbcLa + ITS2) run sequenced on a Flongle flow cell with a MinION Mk1B nanopore sequencer.
4. A medium-scale (292 dipteran specimens), full-length (∼1,900 bp) 18S run sequenced on a Flongle flow cell on a MinION Mk1B nanopore sequencer.

**Table 1:** Experiments used to demonstrate the capacity of BIP to generate barcode data from various types of sequence data, including different amplicon lengths, markers, and sequencing platforms.

| RUN | MARKER | APPROX. AMPLICON LENGTH (BP) | SEQUENCING TECHNOLOGY | SPECIMENS |
|---|---|---|---|---|
| #1 | COI | 658 | Nanopore (ONT) | 100,320 arthropods |
| #2 | COI | 418 | Illumina 2 x 300 PE | 380 arthropods |
| #3 | rbcLa + ITS2 | 550 + 350 | Nanopore (ONT) | 95 flowering plants |
| #4 | 18S | 1900 | Nanopore (ONT) | 292 arthropods |

As these specimens were previously barcoded on a Sequel IIe (Hebert *et al*. 2025), most have known target barcodes and thus possess a ground truth suitable for validating BIP under various analytical conditions. The following text provides a brief description of how each validation dataset was prepared:

#### Validation Run #1

The specimens used for Run #1 were previously barcoded (Hebert *et al*. 2025), and consist of 100,320 Malaise-trapped arthropod specimens representing > 21,000 BINs from 377 families, mostly from five insect orders (76% of specimens were Diptera, 9% Hymenoptera, 5% Coleoptera, 4% Lepidoptera, 4% Hemiptera, and 1.6% from 15 small orders). The original publication provides further details; briefly, the 100,320 specimens were amplified in eleven batches of 9,120 UMI-tagged PCRs. The eleven batches were pooled and tagged with eleven unique Native Barcodes from the ONT kit SQK-NBD114.24. The resulting sequence reads were demultiplexed via the Native Barcodes with MinKNOW into eleven FASTQ files, which were independently analyzed using BIP (employing the same 9,120 unique UMI tags for demultiplexing each time). Analysis proceeded with BIP’s default COI parameters. The results of each of the eleven BIP analyses were combined for comparison to the known BIN composition of the original 100,320 specimens. As a final quality measure, OTUs with fewer than 550 unambiguous bases were discarded.

#### Validation Run #2

The 380 specimens used for Run #2 were from the same project as those used for Run #1 and were selected based on moderate-high sequencing success (> 75%) and spanning multiple arthropod classes (mainly Insecta and Collembola). PCR was performed in two steps, with the first using the primers BF3 + BR2 (Elbrecht *et al*. 2019) which amplify a 418 bp region of COI. Reagent components are provided by Hebert et al. (2018). Thermocycling consisted of enzyme activation at 95°C for 2 min, 30 cycles of [95°C for 30 sec, 46°C for 30 sec, 72°C for 50 sec] and a final extension of 72°C for 5 min. The PCR1 products were used as template for PCR2, which served to add Illumina P5 and P7 adapters to the amplicons. PCR2 reactions were identical in composition to PCR1 and thermocycling consisted of enzyme activation at 94°C for 2 min, 15 cycles of [94°C for 40 sec, 51°C for 1 min, 72°C for 1 min], and a final extension of 72°C for 5 min. The PCR2 products were pooled and purified using magnetic beads; briefly, an equal volume of PCR2 products was pooled and 400 µL of the pool was mixed with 320 µL of Cytiva Sera-Mag beads (Thermo Fisher Scientific). The beads were washed three times with 80% ethanol and eluted in 30 µL of TE buffer (10 mM Tris-HCl, 0.1 mM EDTA, pH 8.0). The purified pool was size-selected for 500–700 bp using a BluePippin (Sage Science) equipped with a 0.25-1.5 kb 1.5% dye-free agarose gel cassette. The size-selected products were purified using magnetic beads as above, this time using a 1:1 DNA:bead ratio and eluting into 15 µL TE. The purified library was quantified using a Qubit 3.0 fluorometer (Thermo Fisher Scientific) and amplicon length distribution was assessed using a 2100 Bioanalyzer (Agilent). The library was then prepared for sequencing on an Illumina P1 flow cell following the manufacturer’s instructions for 2 x 300 bp paired-end (PE) sequencing. Sequencing was performed on a NextSeq 2000 using a P1 XLEAP-SBS 600 cycle kit. The resulting R1 and R2 read sets were randomly subsampled down to 1M reads each and used as input for BIP analysis. As a final quality measure, OTUs with fewer than 200 unambiguous bases were discarded.

#### Validation Run #3

The plant dataset consisted of 95 specimens from 13 flowering plant orders, sequenced simultaneously for a plastid marker (rbcLa) and a nuclear marker (ITS2). DNA was extracted using a silica-membrane based approach (Ivanova, Dewaard, and Hebert 2006). Amplification was performed in 12.5 µL reactions consisting of 6.25 µL of 10% trehalose dihydrate (Fluka Analytical), 1.4 µL of sterile water, 1.25 µL of 10X Platinum Taq buffer (Invitrogen), 0.625 µL of DMSO (Sigma-Aldrich), 0.625 µL of 50 mM MgCl2 (Invitrogen), 0.125 µL of each 10 µM primer, 0.0625 µL of 10 mM dNTP (KAPA Biosystems), 0.060 µL of 5 U/µL Platinum Taq DNA Polymerase (Invitrogen), 0.0125 µL of BSA (Sigma-Aldrich), and 2 µL of DNA. Amplification of rbcLa and ITS2 used the primers rbcLaF + rbcLaR (Tate and Simpson 2003, Kress and Erickson 2007) and ITS-3p62plF1 + ITS-p4R2 (Kolter and Gemeinholzer 2021), respectively, and the following thermocycling conditions: enzyme activation at 94°C for 2 min, 35 cycles of [94°C for 40 sec, 53°C for 40 sec, 72°C for 45 sec], and a final extension of 72°C for 5 min. Following PCR, each reaction was pooled and purified with magnetic beads as above and prepared for sequencing following the manufacturer’s instructions for ONT’s ligation sequencing kit (SQK-LSK114). The prepared library was sequenced on a MinION sequencer equipped with a Flongle flow cell (FLO-FLG114).

#### Validation Run #4

The arthropod 18S dataset consisted of 292 specimens in the fly family, Cecidomyiidae. DNA extraction followed the same methods as Run #1 and full-length (∼1,900 bp) 18S amplification followed a 2-step PCR strategy. All PCR components were the same as Run #1 except for the primers. PCR1 used the primers 18S-CL-F3 + 18S-CL-R1 (Carta and Li 2018) (tailed with universal adapter sequences) and the following thermocycling conditions: enzyme activation at 94°C for 2 min, 5 cycles of [94°C for 40 sec, 45°C for 40 sec, 72°C for 2 min], 40 cycles of [94°C for 40 sec, 51°C for 40 sec, 72°C for 2 min], and a final extension of 72°C for 5 min. The PCR1 products were used as template for PCR2, which served to add UMIs to the amplicons. PCR2 used universal adapter-tailed UMIs as primers and thermocycling consisted of enzyme activation at 94°C for 2 min, 20 cycles of [94°C for 40 sec, 64°C for 1 min, 72°C for 1 min], and a final extension of 72°C for 10 min. The PCR2 products were pooled and purified using magnetic beads. Library preparation followed the manufacturer’s instructions for amplicon sequencing (SQK-LSK114) and sequencing was performed on a MinION sequencer equipped with a Flongle flow cell (FLO-FLG114). The raw reads were used as input for BIP and, as a final quality measure, only OTUs with ≥1,200 (unambiguous) bases were retained.

As Run #1 reflects the most common use case for DNA barcoding (animals barcoded with 658 bp COI), the results generated by BIP were directly compared to those produced by its closest functional analogue, ONTbarcoder 2.3.0 (Srivathsan *et al*. 2024). ONTbarcoder was used in ‘Conventional Barcoding’ mode with default parameters. ONTbarcoder does not match sequences to a reference library, so we used vsearch - -sintax and vsearch --usearch_global to assign taxonomy and match BINs to sequences, respectively. All analyses used the July 2026 COI BOLDistilled library (Prosser *et al*. 2025). ONTbarcoder produced two duplicate barcodes for three specimens in its output (R06_S2046 in R06_output/runsummary.xlsx; R07_S4386 in R07_output/runsummary.xlsx; R08_S5422 in R08_output/runsummary.xlsx), and we removed one of each. We did not compare BIP to NGSpeciesID (Sahlin, Lim, and Prost 2021) because previous work demonstrated that it is outperformed by ONTbarcoder (Srivathsan *et al*. 2021).

### Pipeline Comparisons

We compared sequence results from BIP and ONTbarcoder to those currently on BOLD. The BOLD records represent a ‘gold standard’ for sequence accuracy because they were generated by SMRT sequencing on a Sequel IIe and subsequently curated by experts. Bioinformatic analysis used a desktop computer provisioned with an AMD Ryzen 9 32-core CPU, 64 GB RAM, an NVIDIA GeForce RTX 4090 GPU, running Ubuntu 22.04 LTS. Downstream data analysis of consensus sequences used the following R packages: *readxl* (Wickham and Bryan 2019), *tidyverse* (Wickham *et al*. 2019), and *writexl* (Ooms 2025). Functions in *Biostrings* (Pagès *et al*. 2025) were also widely used to process sequences. Where required, BIP and ONTbarcoder sequences were aligned using `pwalign` (Aboyoun and Gentleman 2025).

## RESULTS

### Run #1: BIP analysis of 100,320 specimens sequenced for COI on ONT

BIP v1.1.0 analyzed 17.6M raw ONT reads generated from sequencing 100,320 arthropod specimens. Approximately 9.5M (54%) of the reads were assigned to an OTU, with most (9M) belonging to what BIP designated as target barcodes (Fig. 2A). BIP identified a target barcode (comprising 21,564 unique BINs) for 91,682 (91%) of the 100,320 specimens. Additionally, BIP identified 13,523 non-target sequences (NTS) (comprising 4,192 unique BINs) from 11,284 specimens. Overall, 82.3% of specimens had only target sequences while 9.1% had both target and NTS (Fig. 2B). A small fraction (2.2%) of specimens had only NTS, while 6.4% of specimens failed to return a sequence. Most specimens (89.8%) that yielded a sequence had a single OTU, though some generated as many as eight OTUs (Fig. 2C). Median read depth was ∼5.5× greater for target barcodes (93) than for NTS (17) (Fig. S1).

**Figure 2:**
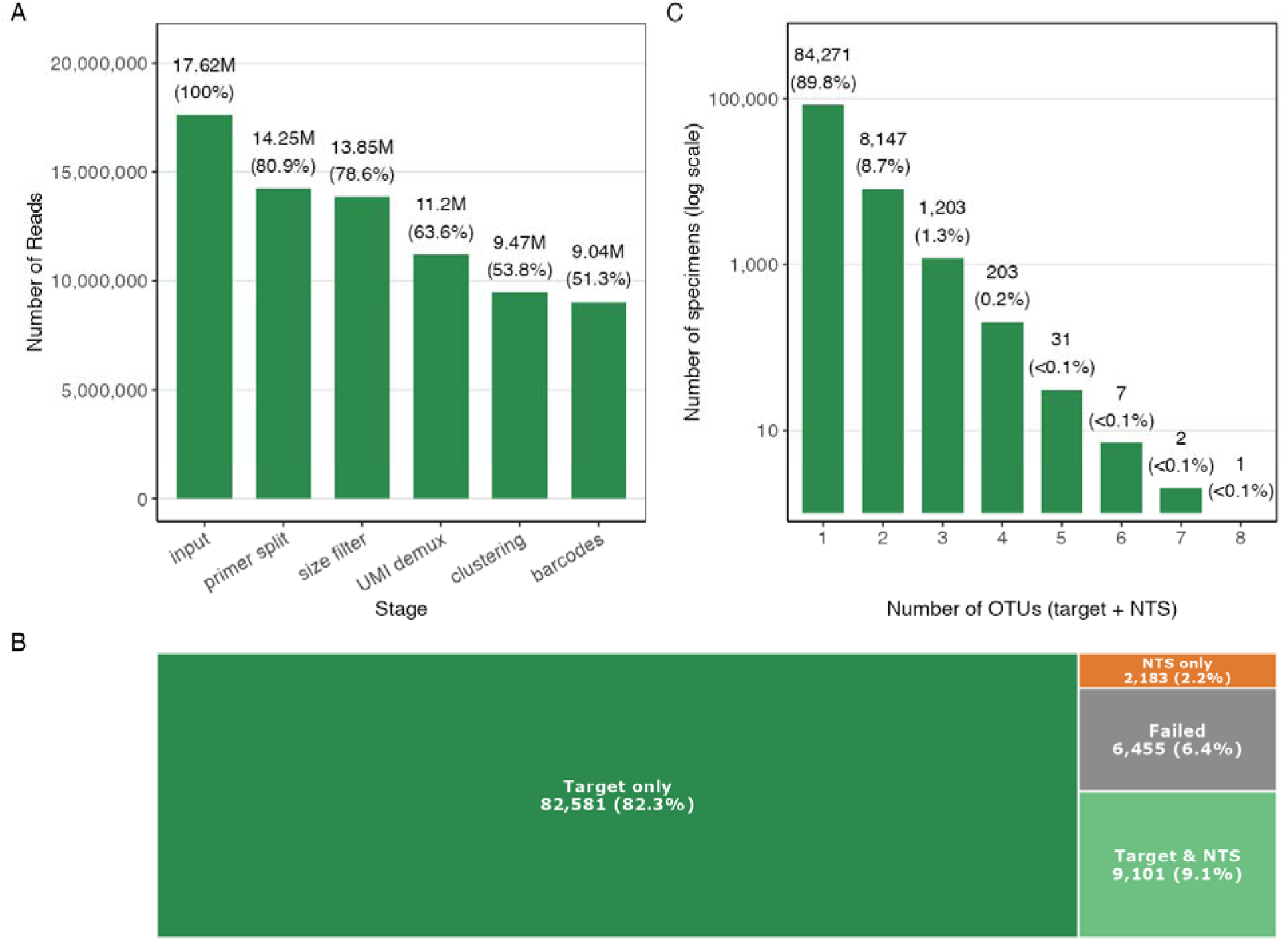
Summary of BIP analysis for validation run #1—100,320 arthropods sequenced for 658 bp COI and run on a MinION Mk1B nanopore sequencer. (A) Read retention throughout the pipeline processing steps. (B) Tree map classifying the BIP results for all 100,320 specimens in the dataset. (C) Number of OTUs per sample in the BIP output for all samples that generated ≥1 OTU (93,865); note the log_10_ scale.

BIP required approximately 6× longer to complete analysis than ONTbarcoder (∼3 h/run vs ∼30 min/run, respectively). The number of reads attributed to each specimen was highly correlated between the methods (*R*^2^ = 0.96; *P* < 0.001; Fig. S2). Across the dataset, 90,806 of 100,320 (90.5%) specimens were assigned a target barcode by both BIP and ONTbarcoder. Sequences were identical for 81,677 (90%) of these specimens, but 2,100 (2.3%) showed sequence divergence exceeding 5% (Fig. 3A). Sequence divergence between BIP target barcodes and ONTbarcoder barcodes was greater for specimens where BIP reported the presence of NTS (Fig. 3B; Fig. S3).

**Figure 3:**
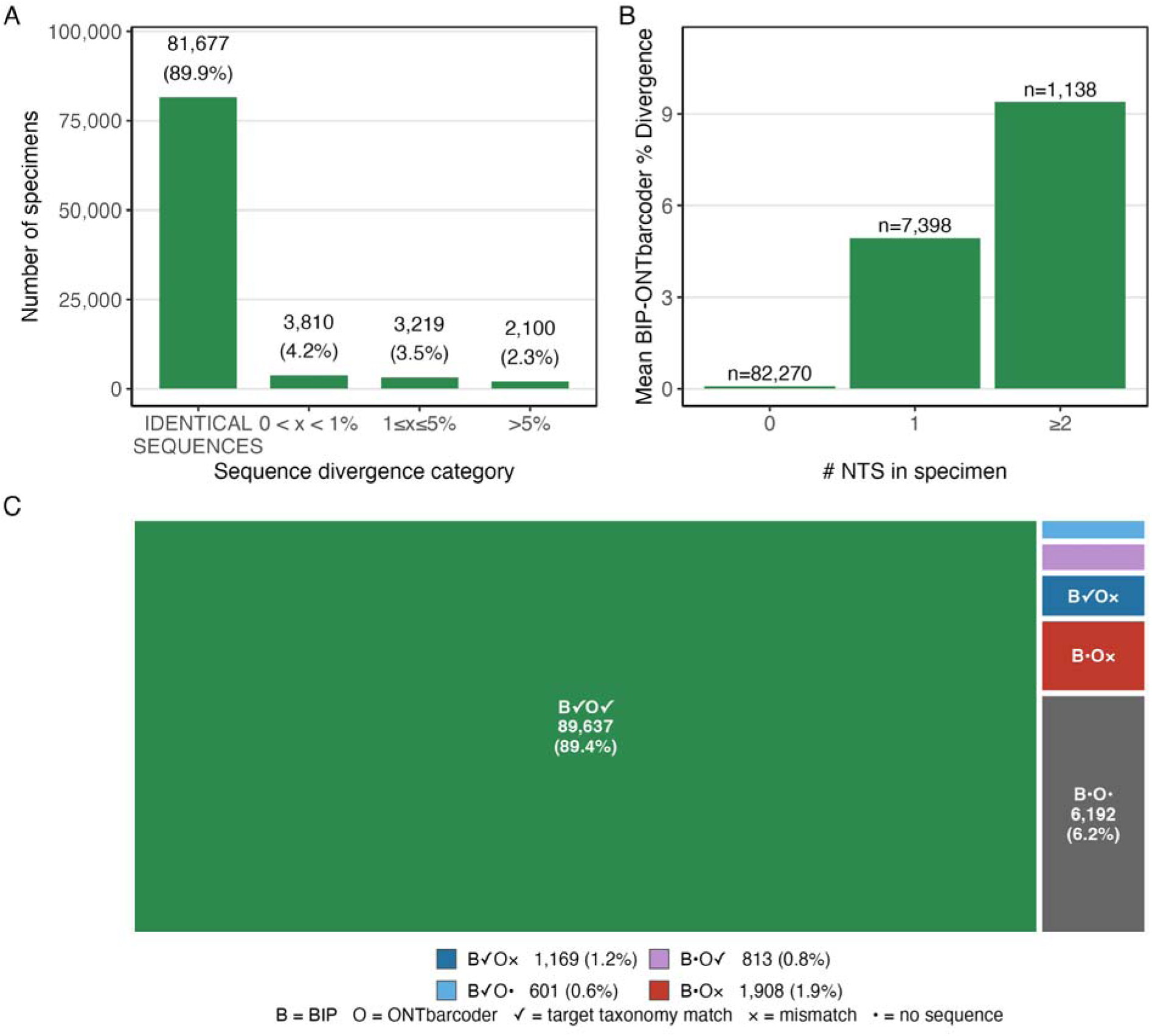
Comparison of BIP target sequences with ONTbarcoder output. (A) Most ONTbarcoder sequences (89.9%) are identical to BIP target sequences, but the others show divergence. (B) Divergence between ONTbarcoder and BIP target barcodes is greater when BIP identifies more NTS. (C) Tree map showing comparative results for each specimen analyzed by BIP and ONTbarcoder. A & B only contain data for the 90,806 specimens for which both BIP and ONTbarcoder generated a barcode; Panel C reflects outcomes for all 100,320 specimens. Note: B× results are not possible because BIP, by definition, matches target taxonomy.

We evaluated the correspondence of barcodes to expected taxonomy using sintax (Fig. 3C). Among all 100,320 specimens, 89.4% gained a (target) barcode meeting the taxonomic match criteria via both pipelines, while 6.2% failed to gain a valid barcode via both pipelines. For 1.9% of specimens, ONTbarcoder returned sequences that did not belong to the target taxon while BIP returned no target barcode. For 0.8% of specimens, ONTbarcoder returned correct barcodes while BIP returned no target; the reverse (a correct barcode from BIP, none from ONTbarcoder) occurred for 0.6% of specimens. (Note that BIP will not return a taxonomically invalid barcode.) Finally, 1.2% of specimens had a correct barcode from BIP and an incorrect barcode from ONTbarcoder. Thus, outcomes differed between BIP and ONTbarcoder for 4.5% of specimens, and only 0.8% of specimens had a better outcome from ONTbarcoder.

Incongruence between BIP and ONTbarcoder was primarily linked to the presence of NTS. Across the dataset, BIP target vs. ONTbarcoder barcode divergence was strongly positively associated with the fraction of a specimen’s reads that BIP attributed to NTS (ρ = 0.63; *P* < 0.001; Fig. 4A; analysis excludes identical sequences). Divergence was greater when the BIP target and ONTbarcoder barcodes were more distant taxonomically. Of the 93,598 barcodes returned by ONTbarcoder, 91.3% match the BIP target barcode within 1%; 3.1% match an NTS within 1%; and 5.5% are more than 1% divergent from both BIP target and NTS (Fig. 4B). In addition, although 94.4% of BIP’s target sequences were 658 bp, 5.6% ranged in length from 640–673 bp, while 99.99% of ONTbarcoder’s sequences were 658 bp (Fig. 4C). Finally, when compared to PacBio Sequel IIe data, BIP outperformed ONTbarcoder: pairwise divergence from Sequel IIe data downloaded from BOLD was considerably lower for BIP than ONTbarcoder (Fig. 4D, Table 2).

**Figure 4:**
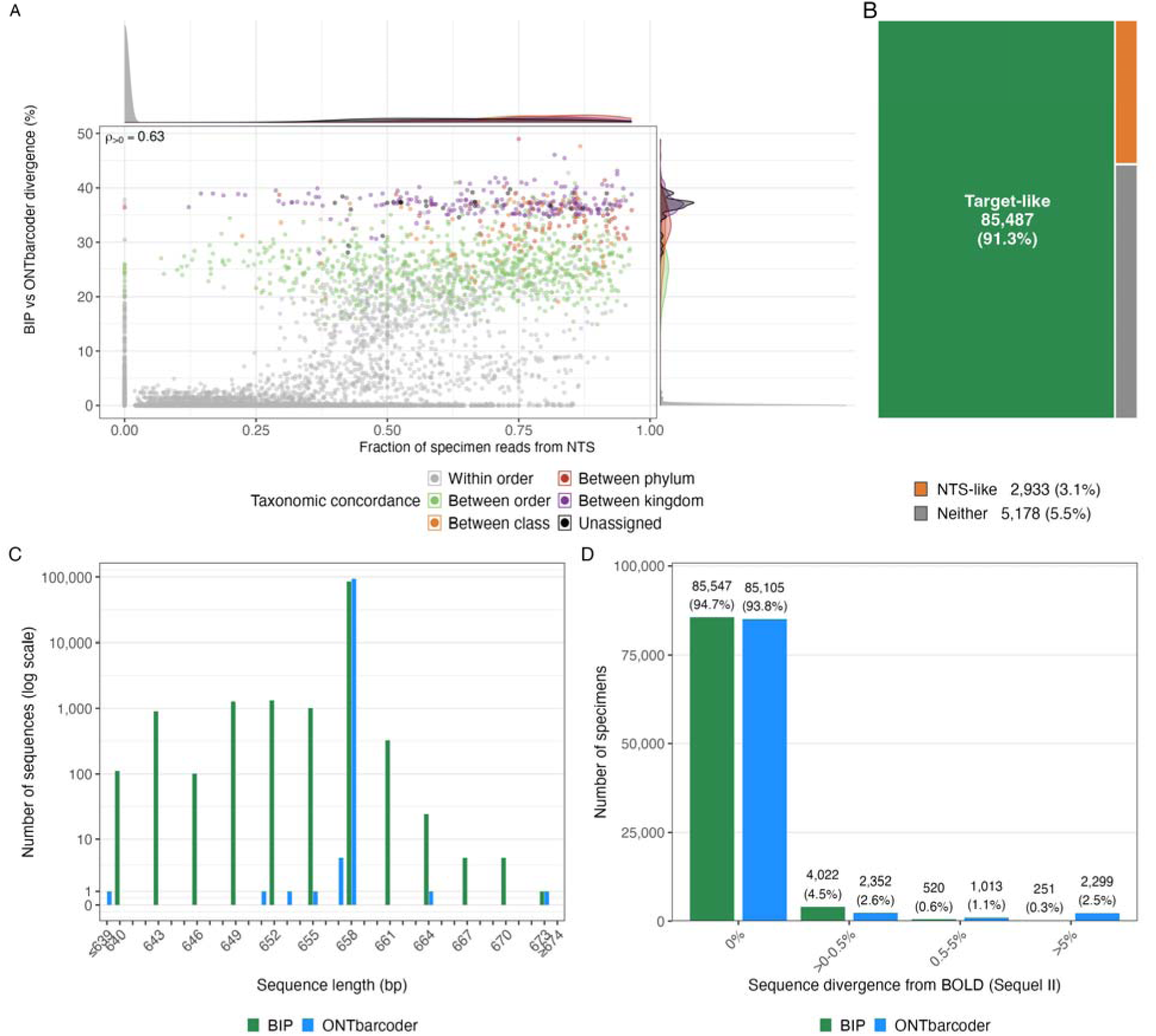
Characteristics of ONTbarcoder barcodes relative to BIP output. (A) Specimens with a higher fraction of reads from NTS in BIP show greater barcode divergence from ONTbarcoder sequences (Spearman’s ρ = 0.63). Density plots at the margins show the distribution of data. (B) Breakdown of all 93,598 ONTbarcoder sequences by whether they were target-like (< 1% divergence from BIP target), NTS-like (> 1% divergence from target & < 1% divergence from a BIP NTS) or neither. (C) Comparison of sequence length distribution between BIP and ONTbarcoder (OBC). Note the pseudo log_10_-scale which allows for the visualization of ‘1’ values. (D) Categorical comparison of sequence divergence of BIP and ONTbarcoder (OBC) to curated PacBio Sequel IIe data downloaded from BOLD.

**Table 2:**
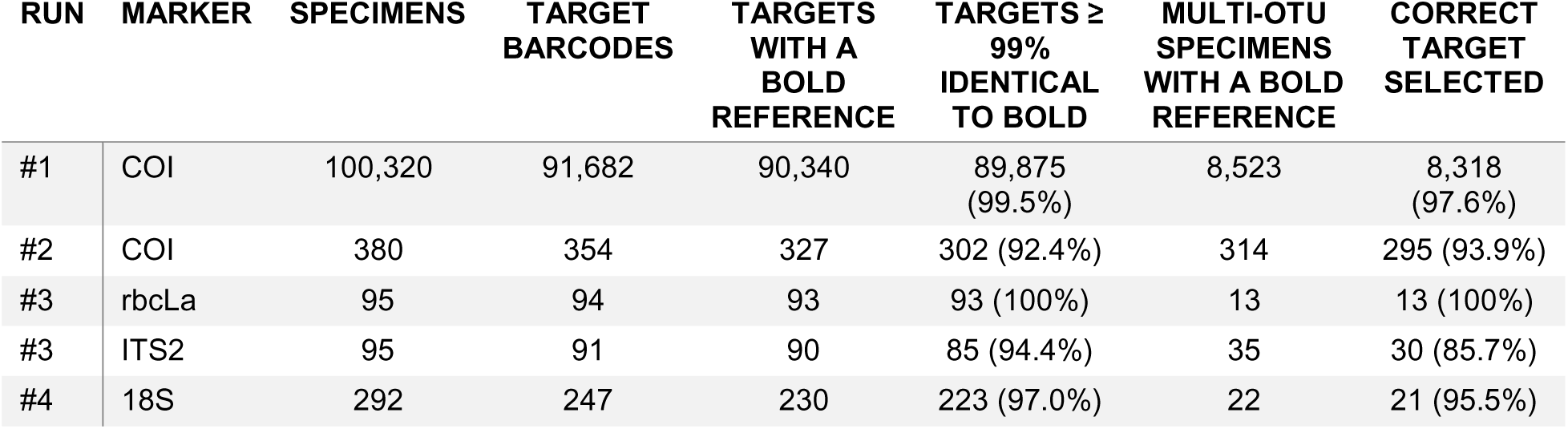
Congruence of BIP target barcodes with reference sequences on BOLD for the four validation runs.

| RUN | MARKER | SPECIMENS | TARGET BARCODES | TARGETS WITH A BOLD REFERENCE | TARGETS ≥ 99% IDENTICAL TO BOLD | MULTI-OTU SPECIMENS WITH A BOLD REFERENCE | CORRECT TARGET SELECTED |
|---|---|---|---|---|---|---|---|
| #1 | COI | 100,320 | 91,682 | 90,340 | 89,875 (99.5%) | 8,523 | 8,318 (97.6%) |
| #2 | COI | 380 | 354 | 327 | 302 (92.4%) | 314 | 295 (93.9%) |
| #3 | rbcLa | 95 | 94 | 93 | 93 (100%) | 13 | 13 (100%) |
| #3 | ITS2 | 95 | 91 | 90 | 85 (94.4%) | 35 | 30 (85.7%) |
| #4 | 18S | 292 | 247 | 230 | 223 (97.0%) | 22 | 21 (95.5%) |

### Run #2: BIP analysis of 380 specimens sequenced for COI on Illumina

The Illumina run produced 132.5M paired-end (PE) reads which were subsampled down to 1M reads—ample depth for 380 specimens. Of those 1M read pairs, 861,171 were successfully merged. After filtering, demultiplexing, and primer trimming, 781,948 (78.2%) of the reads were assigned to an OTU (Fig. 5A). Of the 2,840 OTUs, 354 (12.5%) were designated by BIP to be target barcode sequences while 2,486 (87.5%) were NTS (Fig. 5C). A target barcode was recovered from 354 (93.2%) of the 380 specimens. In contrast to the ONT run, very few specimens (13, 3.4%) yielded only a target barcode. Instead, most specimens (341, 89.7%) yielded both a target and NTS, while 24 (6.3%) yielded only NTS and 2 (0.5%) failed to yield any OTUs (Fig. 5B). When compared to their BOLD counterparts, 302 (92.4%) of the 327 Illumina-generated target barcodes matched with ≥ 99% identity. When a specimen yielded more than one OTU, BIP selected the correct barcode sequence 93.9% of the time (Table 2).

**Figure 5:**
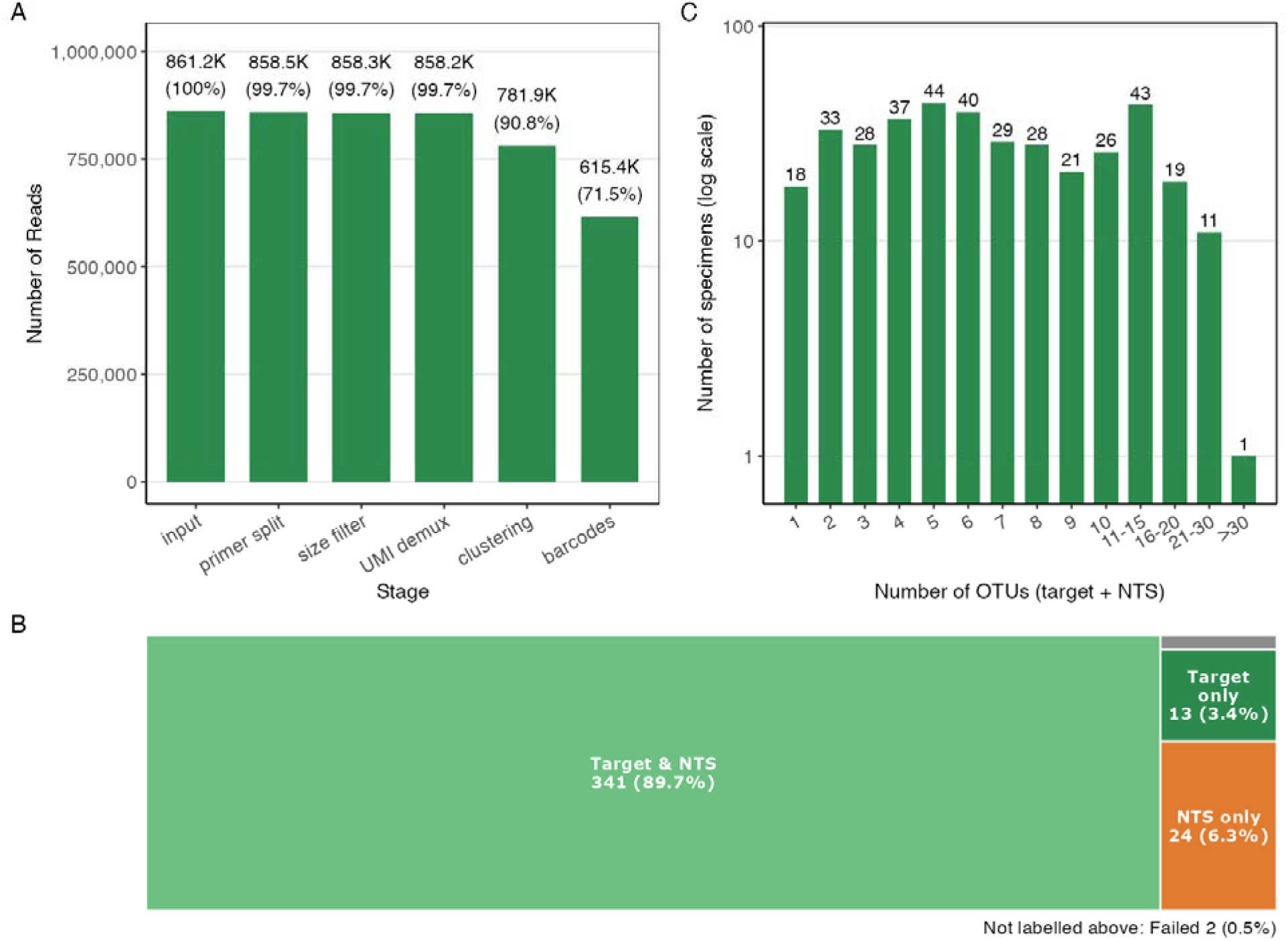
Summary of BIP analysis for validation run #2—380 arthropods sequenced for 418 bp COI on an Illumina NextSeq 2000. (A) Read retention throughout the pipeline processing steps. (B) Tree map classifying the BIP results for all 380 specimens in the dataset. (C) Number of OTUs per sample in the BIP output for all samples that generated ≥1 OTU (378); note the log_10_ scale.

### Run #3: BIP analysis of multiple plant markers (rbcLa + ITS2) on ONT

The multi-marker plant run yielded 979,043 reads with 222,219 (22.7%) belonging to rbcLa and 482,464 (49.3%) belonging to ITS2 (Fig. 6A; Fig. 7A). BIP assigned 60,542 (27.2%) and 58,409 (12.1%) of the rbcLa and ITS2 reads to OTUs, respectively (Fig. 6C; Fig. 7C), and identified an rbcLa target barcode sequence for 94 specimens and an ITS2 target barcode for 91 specimens (Fig. 6B; Fig. 7B). The nuclear ITS2 marker yielded more NTS (62 OTUs) than the plastid rbcLa marker (14 OTUs). When compared to their counterparts on BOLD (93 and 90 records for rbcLa and ITS2, respectively), 93 (100%) of the rbcLa barcodes and 85 (94.4%) of the ITS2 barcodes matched with ≥ 99% identity (Table 2).

**Figure 6:**
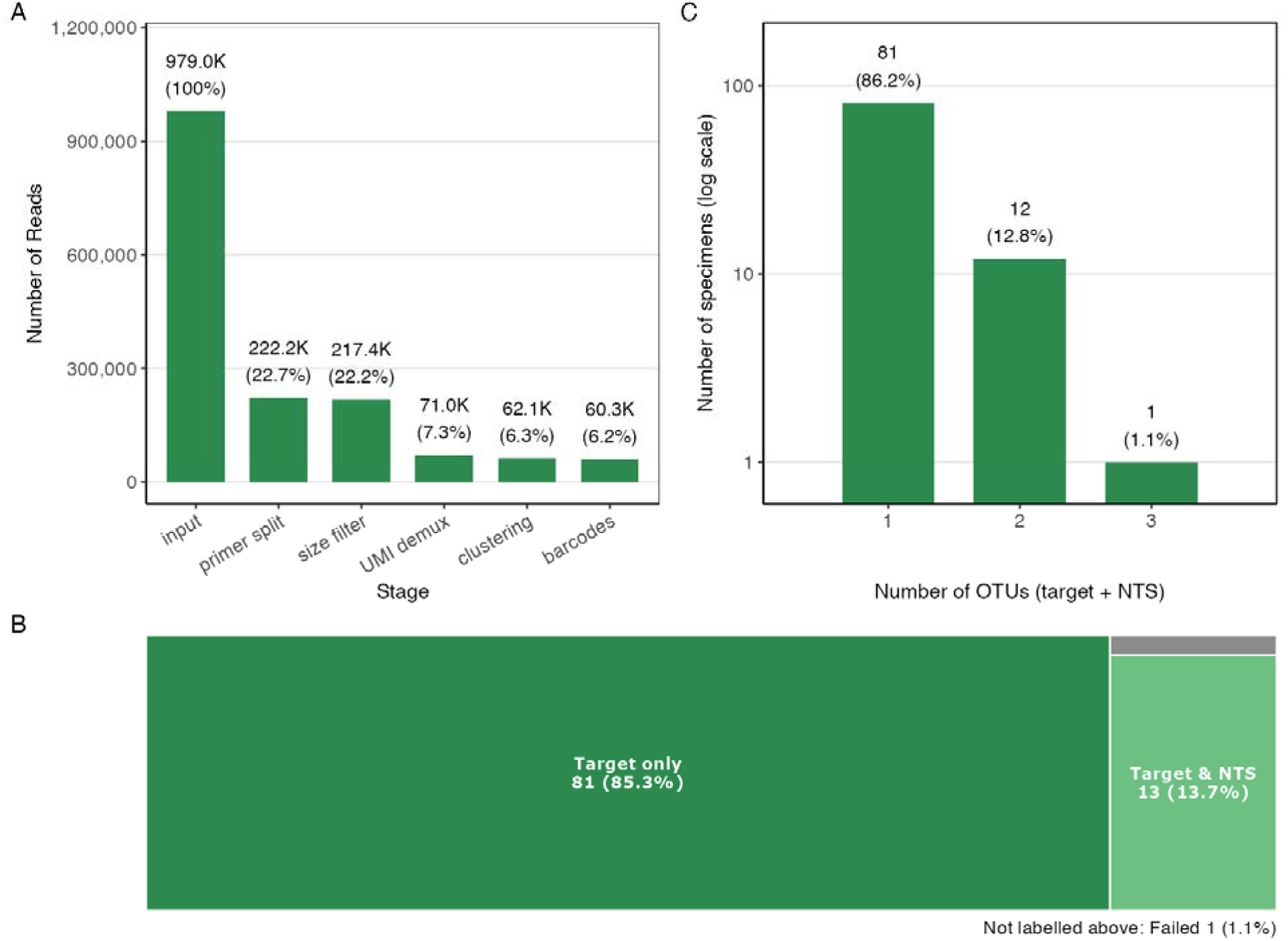
Summary of BIP analysis for validation run #3—95 flowering plants sequenced for ∼550 bp rbcLa on a MinION Mk1B nanopore sequencer. (A) Read retention throughout the pipeline processing steps. (B) Tree map classifying the BIP results for all 95 specimens in the dataset. (C) Number of OTUs per sample in the BIP output for all samples that generated ≥1 OTU (94); note the log_10_ scale.

**Figure 7:**
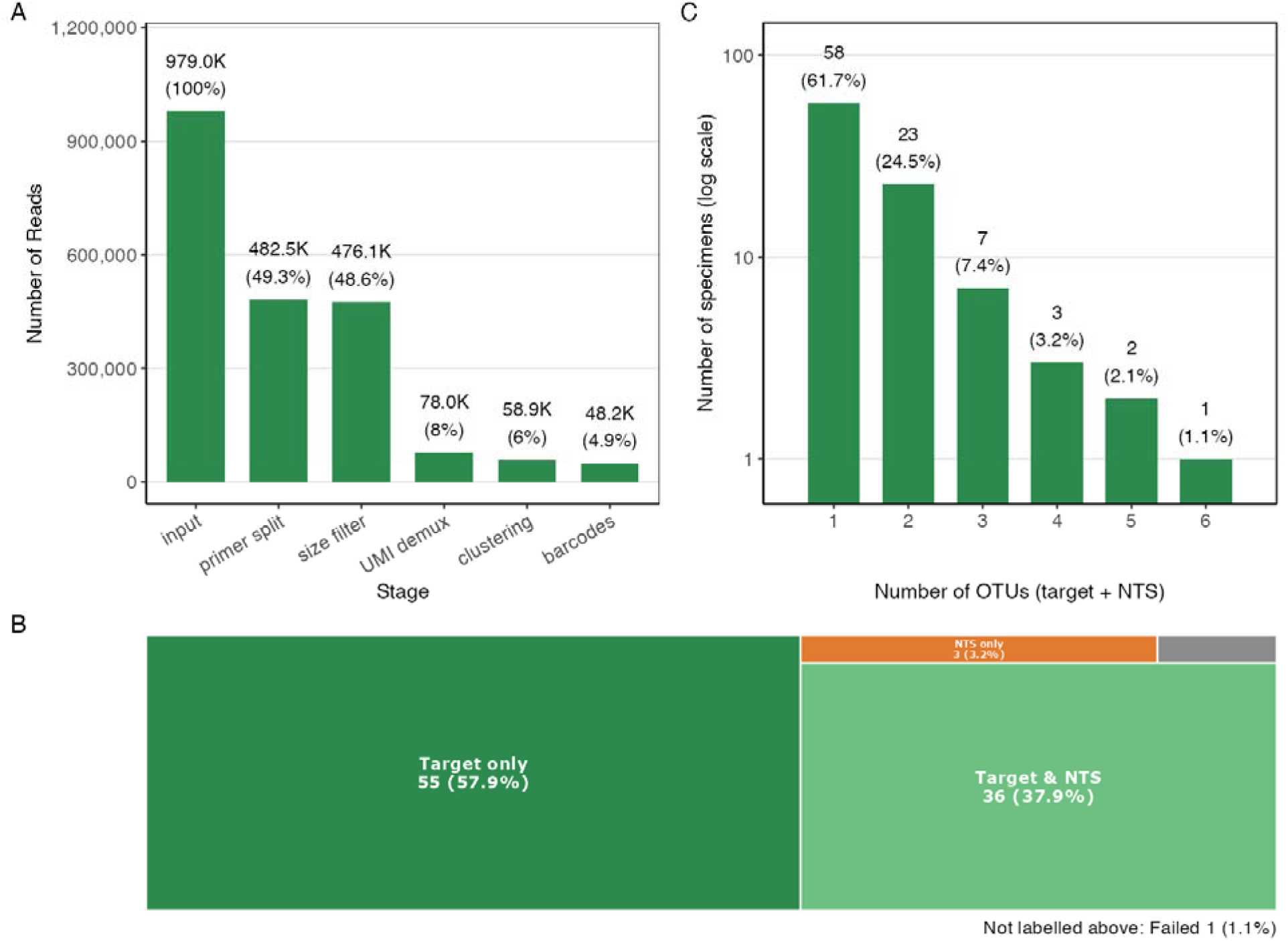
Summary of BIP analysis for validation run #3—95 flowering plants sequenced for ∼350 bp ITS2 on a MinION Mk1B nanopore sequencer. (A) Read retention throughout the pipeline processing steps. (B) Tree map classifying the BIP results for all 95 specimens in the dataset. (C) Number of OTUs per sample in the BIP output for all samples that generated ≥1 OTU (94); note the log_10_ scale.

### Run #4: BIP analysis of the full-length nuclear 18S marker on ONT

The 18S run generated 1,674,733 reads, of which only 32,921 (2.0%) were assigned to OTUs because most reads were shorter than the expected minimum length of 1,200 bp (Fig. 8A). Inspection of the short reads revealed them to be spurious, random (i.e., non-18S) amplicons, indicating non-specific primer binding during PCR. Nevertheless, the reads derived from 18S sufficed to generate target 18S barcodes for 247 (84.6%) of the 292 specimens (Fig. 8B); 25 NTS were also recovered from 23 specimens (Fig. 8C). Of the 247 target barcodes produced by BIP, 230 had counterparts on BOLD, of which 223 (97.0%) matched their BOLD counterparts with ≥ 99% identity.

**Figure 8:**
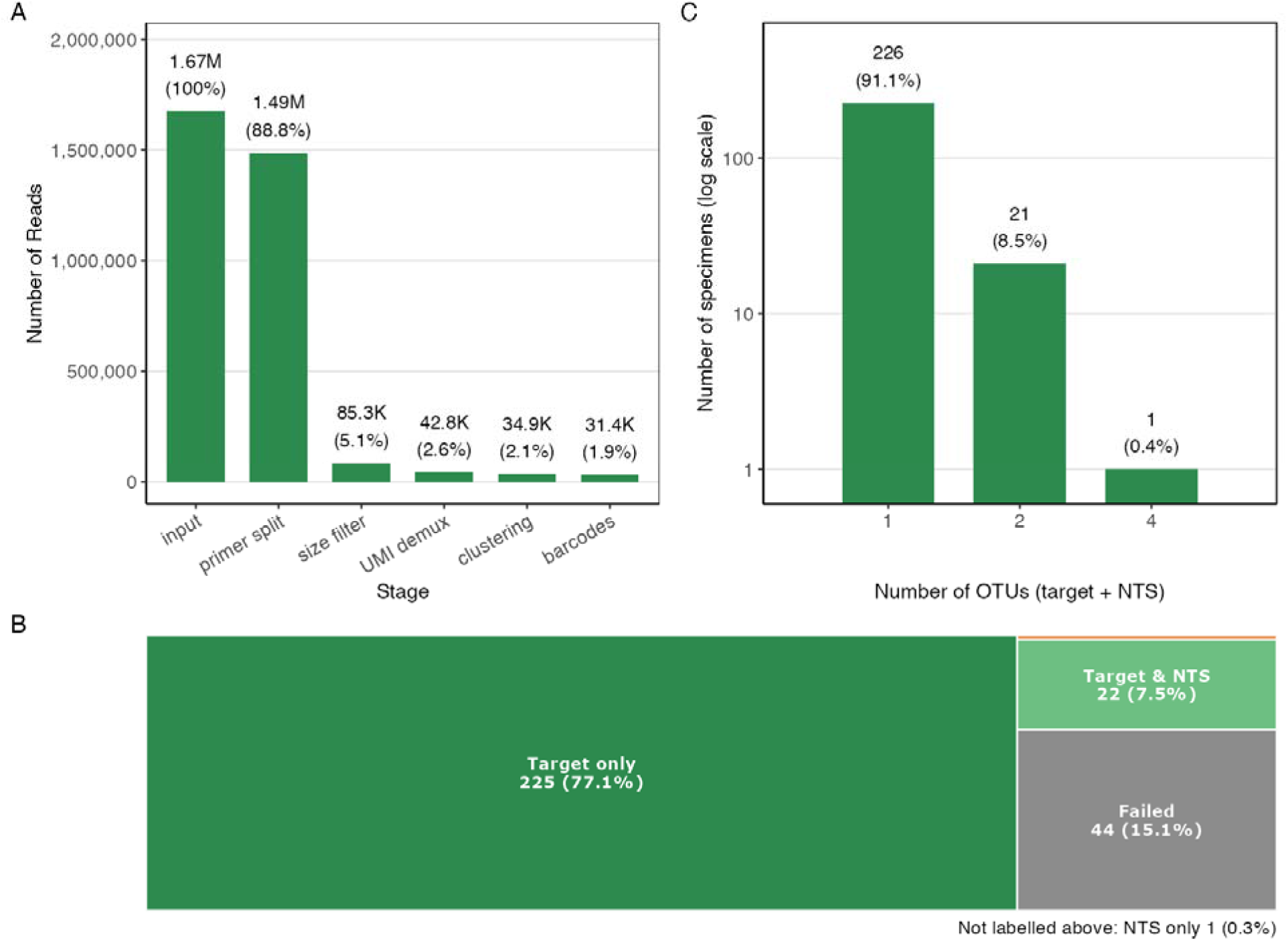
Summary of BIP analysis for validation run #4—292 dipterans sequenced for ∼1,900 bp 18S on a MinION Mk1B nanopore sequencer. (A) Read retention throughout the pipeline processing steps. (B) Tree map classifying the BIP results for all 292 specimens in the dataset. (C) Number of OTUs per sample in the BIP output for all samples that generated ≥1 OTU (248); note the log_10_ scale.

## DISCUSSION

Although many pipelines can generate OTUs from multiplexed sequence data (Hakimzadeh *et al*. 2024), only BIP and ONTbarcoder are specifically designed to support DNA barcode analysis. The two differ in one key respect: ONTbarcoder presumes that the sequences recovered from a specimen represent amplicons derived solely from the target gene region in this specimen. Consequently, although ONTbarcoder demultiplexes all reads associated with a specimen, it generates a single consensus sequence from them. As a result, if DNA sequences from more than one species are present, ONTbarcoder either uses the dominant haplotype or merges the sequences into one consensus, inserting Ns at sites with a substitution. It therefore cannot protect the barcode from complications introduced by multiple templates in a DNA extract, whether they arise from contamination, NUMTs, endosymbionts, parasites, or parasitoids.

The present analysis examined data from two sequencing platforms, four markers, three genomes (mitochondrial, chloroplast, nuclear), and five primer sets. When more than one OTU per specimen was present, the degree to which BIP inferred the correct barcode (i.e., the sequence currently on BOLD) depended on the marker. BIP selected the correct target OTU > 90% of the time for all COI analyses, 100% for rbcLa, > 95% for 18S, and > 85% for ITS2 (Table 2). The lower success rate for ITS2 is explained by its high intra-specimen sequence variation. For example, in those cases where the target ITS2 barcode selected by BIP differed from that on BOLD, the BOLD sequence matched an NTS from BIP that was, in every case, the most abundant OTU. BIP bypassed these OTUs because its tie-breaking algorithm prioritizes taxonomic agreement and expected sequence length over read depth. For length-variable, intra-genomically diverse markers such as ITS2, prioritizing read depth in tie-breaking would improve target selection. Indeed, read depth prioritization resulted in a substantial improvement in the congruence between BIP’s target barcodes and BOLD for ITS2 (from 85% to 100%) but not for COI, rbcLa, or 18S (Table S1).

We carefully compared results generated by BIP with those from ONTbarcoder. Although both pipelines analyzed most sequences correctly, BIP was more accurate as 3% of the barcodes returned by ONTbarcoder did not originate from the target specimen. In other cases where specimens possessed both target and non-target DNA, ONTbarcoder usually produced a sequence which diverged from that generated by BIP. While BIP and ONTbarcoder both returned valid barcodes for approximately 90% of specimens, ONTbarcoder generated taxonomically incorrect barcodes for 1.9% of specimens that BIP classified correctly as NTS. BIP also delivered barcode sequences that reflect real length variation in the COI amplicon arising as a consequence of insertions and deletions (Fig. S4), as commonly occurs in haplodiploid organisms (Pakrashi, Thompson, and Hebert 2025). Finally, BIP delivered non-target sequences from 11% of the specimens analyzed for COI. While these sequences must be inspected carefully to ascertain their source (see Supplemental Text), some provide important biological insights (e.g., host-parasitoid interactions).

BIP is the bioinformatics pipeline used to recover DNA barcodes from millions of specimens each year at the Centre for Biodiversity Genomics (CBG). This level of production is only possible because of its ability to ingest raw sequence data and output target barcode sequences without the need for pre- or post-processing. The present publication provides transparency to researchers using CBG-generated data and also allows them to independently reproduce results and validate barcode data. Furthermore, by providing access to both target and non-target sequences, BIP provides a basis for gaining insights into species interactions (Dovydaitis, Schmidt, and Gerth 2026). This information is latent within every barcode dataset generated by next-generation sequencing and BIP helps to illuminate it.

## AUTHOR CONTRIBUTIONS

- SP: Software design, initial build, data analysis, manuscript revision
- EO: Software design advice
- NB: Software containerization and refinement
- KT: Manuscript first draft, benchmarking, data analysis, manuscript revision
- RF: Software design (error correction), sequence acquisition, manuscript revision
- PH: Project supervision, manuscript structure and revision, resource acquisition

## ACKNOWLEDGEMENTS

We thank CBG staff for overseeing DNA extraction and PCR for the “100K dataset”. Evgeny Zakharov, Dirk Steinke, Renee Miskie, and Jayme Sones provided useful feedback on BIP’s design. Masha Kuzmina provided the plant DNA. Daniel Parsons and Jordan Beasley provided feedback. This study was enabled by support to PDNH from the Government of Canada through Genome Canada and Ontario Genomics (OGI-208 and OGI-233), and by the New Frontiers in Research Fund (NFRFT-2020-00073). Infrastructure required to carry out the work was acquired with support from the Canada Foundation for Innovation (CFI) and the Gordon and Betty Moore Foundation. A Major Science Infrastructure award (MSI 42450) from CFI plays a key role in sustaining the CBG’s informatics and sequencing platforms. Analysis code was written with the assistance of Claude Code.

## DATA ACCESSIBILITY

All original raw data and analysis code are available on FigShare: https://doi.org/10.6084/m9.figshare.33023282.

## SUPPLEMENTARY MATERIALS

### Supplemental Text

By recovering both target and non-target sequences, BIP provides the opportunity to gain insights into species interactions. For example, in the present study, it revealed that 274 of the 100,320 specimens in Run #1 were parasitized by a nematode (Supplemental File S1). Similar information is latent within every barcode dataset generated by next-generation sequencing but the results must be interpreted with care.

Although detecting template diversity is critical for gaining insights into species interactions, it needs emphasis that many NTS do not represent a meaningful biological association (e.g., those arising through contamination or tag jumps). For a co-occurrence to be considered as a biological association, the following criteria should be considered:

**1) The association must be biologically realistic**. For example, life history, known behavior, geographical range, and other biological factors can render a potential association very unlikely.
**2) The association should be supported by several independent detections**. Although a single co-occurrence may be coincidental, the repeated detection of sequences from two species is likely to reflect a genuine biological association.
**3) The association should be supported by high-quality sequence data**. The association should be supported by NTS that are free of errors and, if protein-coding, should not contain frameshift causing indels or stop codons. Ideally, they should also have a high read depth. NTS supported by read counts similar to those in negative controls are likely to reflect cross-contamination.

### Supplemental Tables and Figures

**Table S1:**
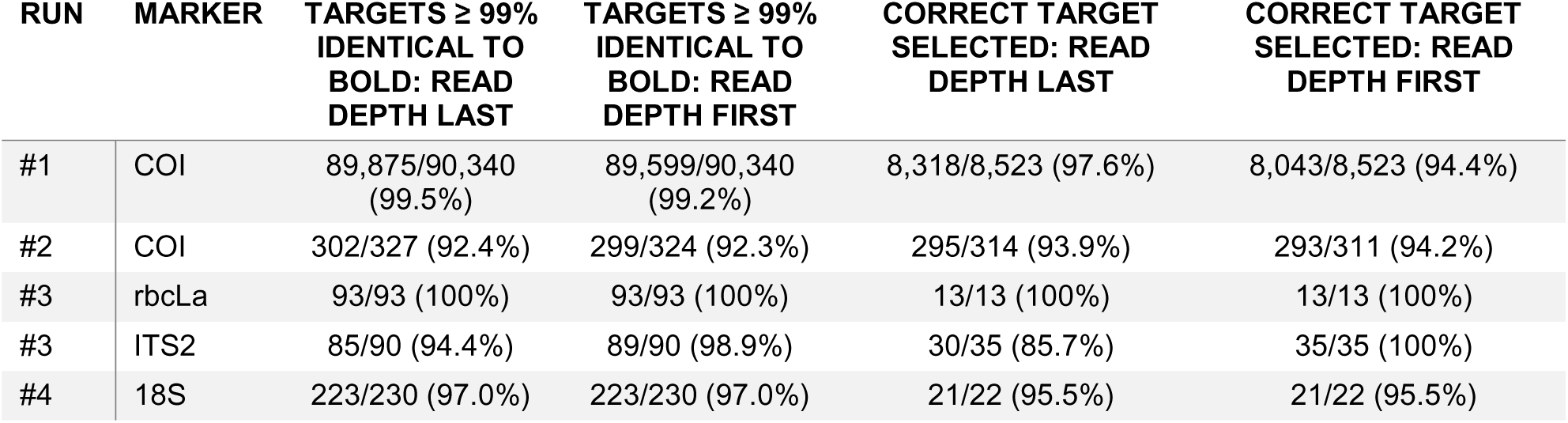
Congruence of BIP target barcodes with reference sequences on BOLD when target selection prioritizes read depth for tie-breaking (read depth > lower-rank taxonomic match > ambiguous bases > distance from the expected amplicon length).

| RUN | MARKER | TARGETS ≥ 99% IDENTICAL TO BOLD: READ DEPTH LAST | TARGETS ≥ 99% IDENTICAL TO BOLD: READ DEPTH FIRST | CORRECT TARGET SELECTED: READ DEPTH LAST | CORRECT TARGET SELECTED: READ DEPTH FIRST |
|---|---|---|---|---|---|
| #1 | COI | 89,875/90,340 (99.5%) | 89,599/90,340 (99.2%) | 8,318/8,523 (97.6%) | 8,043/8,523 (94.4%) |
| #2 | COI | 302/327 (92.4%) | 299/324 (92.3%) | 295/314 (93.9%) | 293/311 (94.2%) |
| #3 | rbcLa | 93/93 (100%) | 93/93 (100%) | 13/13 (100%) | 13/13 (100%) |
| #3 | ITS2 | 85/90 (94.4%) | 89/90 (98.9%) | 30/35 (85.7%) | 35/35 (100%) |
| #4 | 18S | 223/230 (97.0%) | 223/230 (97.0%) | 21/22 (95.5%) | 21/22 (95.5%) |

**Figure S1:**
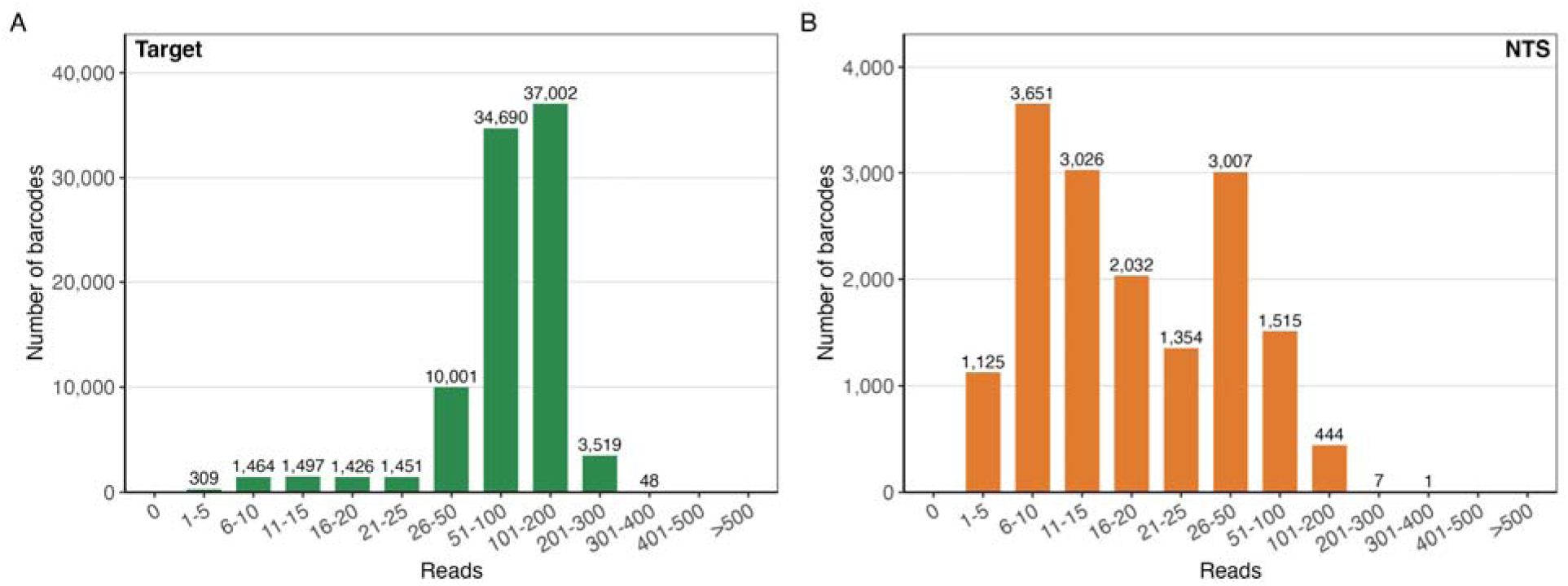
Read depth of OTUs in validation run #1 (100,320 arthropods) analyzed by BIP v1.1.0. (A) Target sequences. (B) NTS. Note the difference in *y*-axis scales.

**Figure S2:**
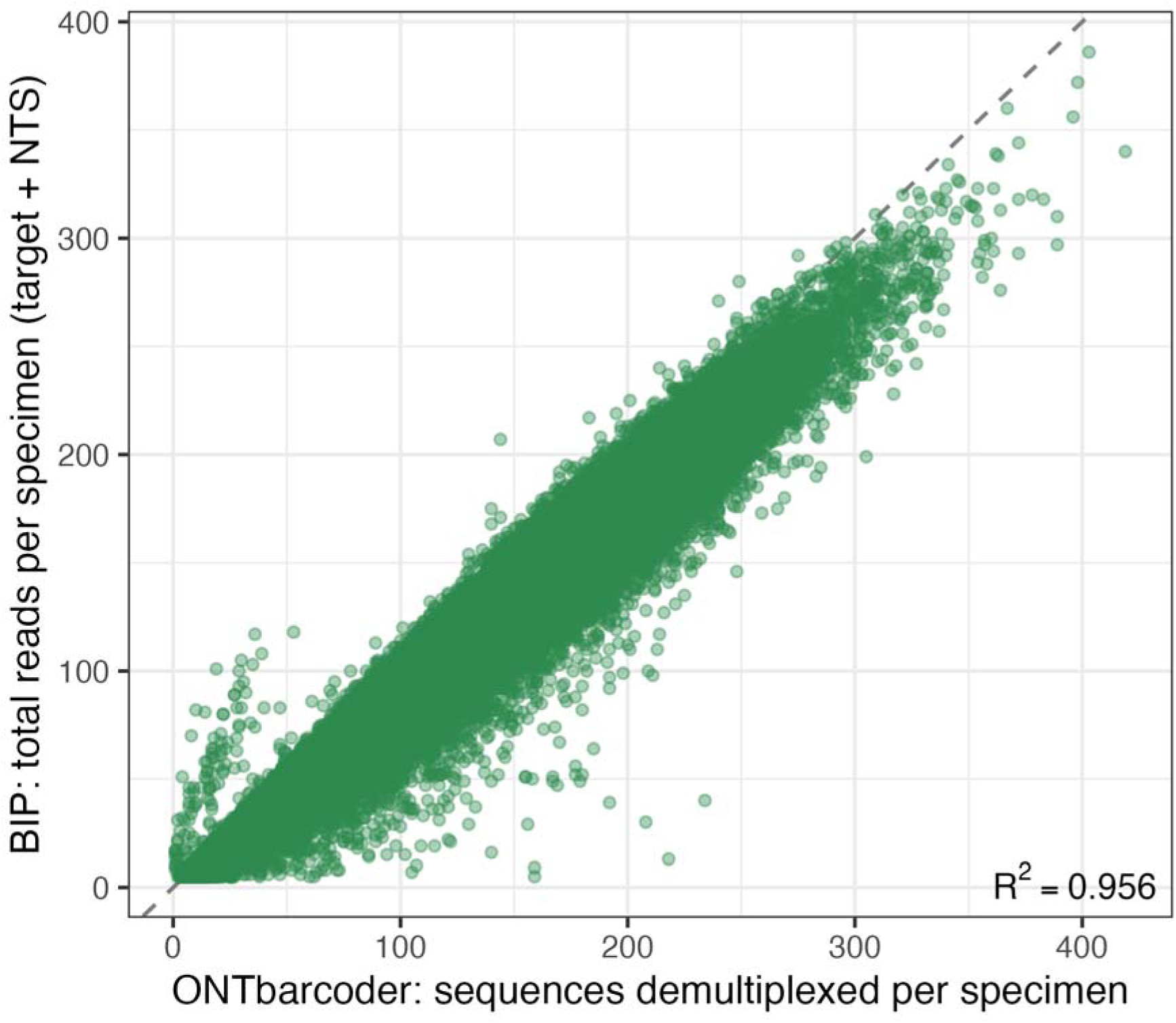
Correlation between sequence depth from ONTbarcoder and BIP. The ‘demultiplexed reads’ value is reported by ONTbarcoder, while BIP reports the total number of reads in target barcodes and NTS.

**Figure S3:**
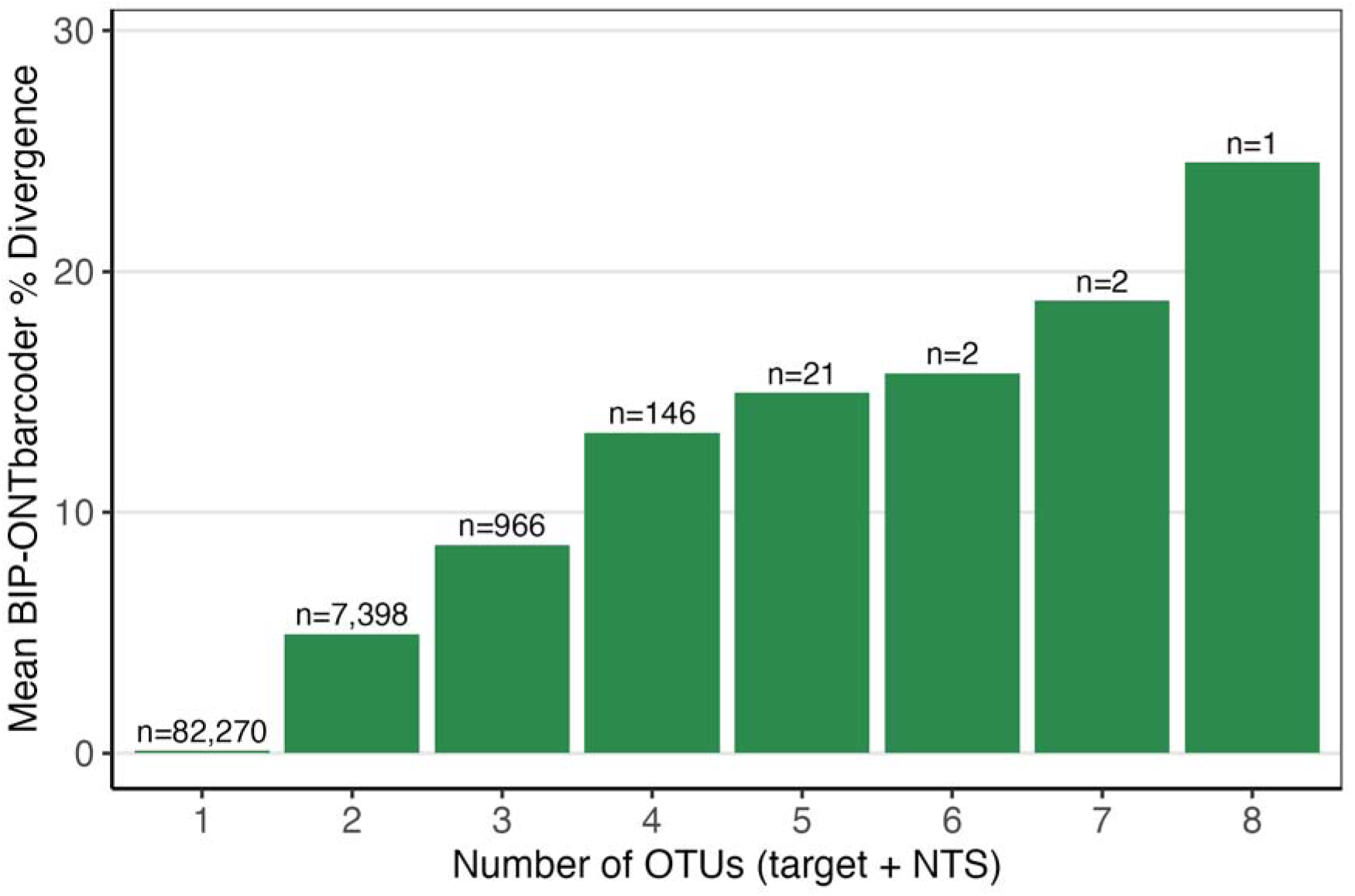
Data presented in. Fig. 3B **with the ‘**≥ **2’ category displayed fully.** The numbers differ from Fig. 2C because those data reflect BIP output alone. Note the x-axis here shows the number of OTUs, not just NTS.

**Figure S4:**
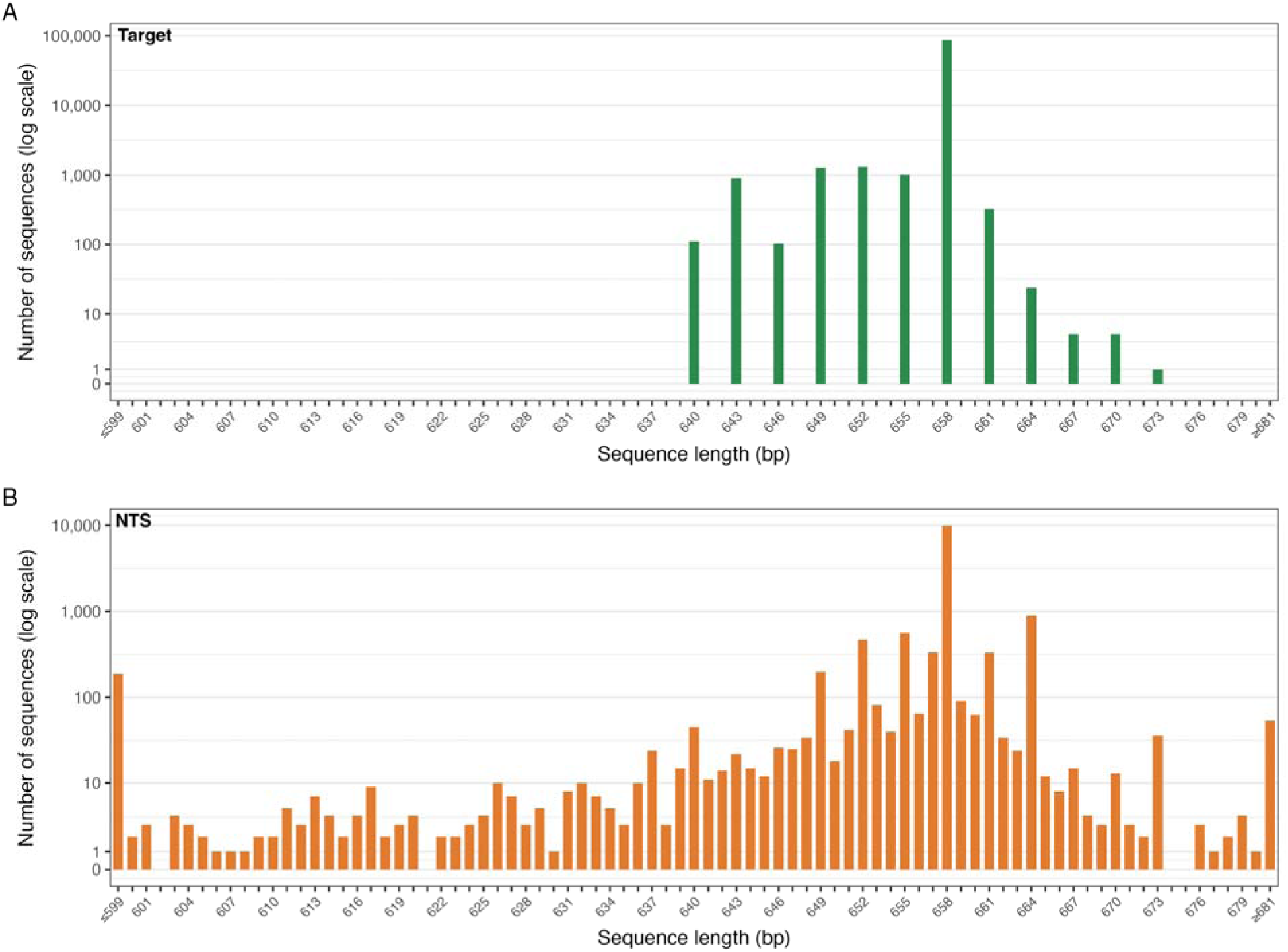
Length of COI sequences for OTUs in validation run #1 (100,320 arthropods) analyzed by BIP v1.1.0. (A) Length variation among COI amplicons in target sequences reflecting 3 bp indels. (B) Length variation among COI amplicons in NTS with the continuous distribution indicating frameshift and non-frameshift indels. Note the difference in *y*-axis scales, which are both on a pseudo-log_10_ scale.

## REFERENCES

Aboyoun P, Gentleman R. Pwalign: Perform Pairwise Sequence Alignments, version 1.0.0. 2025. https://bioconductor.org/packages/pwalign.

Baker CCM, Bittleston LS, Sanders JG et al. Dissecting host-associated communities with DNA barcodes. Philos Trans R Soc Lond B Biol Sci 2016;371(1702):20150328.

Callahan BJ, McMurdie PJ, Rosen MJ et al. DADA2: High-resolution sample inference from Illumina amplicon data. Nat Methods 2016;13(7):581–3.

Carta LK, Li S. Improved 18S small subunit rDNA primers for problematic nematode amplification. Journal of Nematology 2018;50(4):533–42.

Dovydaitis E, Schmidt C, Gerth M. Symbiont interactions bias measures of arthropod biodiversity. EcoEvoRxiv 2026:13318.

Edgar RC, Haas BJ, Clemente JC et al. UCHIME improves sensitivity and speed of chimera detection. Bioinformatics 2011;27(16):2194–200.

Elbrecht V, Braukmann TWA, Ivanova NV et al. Validation of COI metabarcoding primers for terrestrial arthropods. PeerJ 15 July 2019:e7745.

Furneaux B, Anslan S, Somervuo P et al. OptimOTU: Taxonomically aware OTU clustering with optimized thresholds and a bioinformatics workflow for metabarcoding data. In: arXiv 2502.10350. 14 Feb. 2025. arXiv. 10.48550/arXiv.2502.10350.

Hakimzadeh A, Abdala Asbun A, Albanese D et al. A pile of pipelines: An overview of the bioinformatics software for metabarcoding data analyses. Mol Ecol Resour 2024;24(5):e13847.

Hebert PDN, Braukmann TWA, Prosser SWJ et al. A Sequel to Sanger: amplicon sequencing that scales. BMC Genomics 2018;19(1):219.

Hebert PDN, Cywinska A, Ball SL et al. Biological identifications through DNA barcodes. Proc Biol Sci 2003;270(1512):313–21.

Hebert PDN, Floyd R, Jafarpour S et al. Barcode 100K specimens: In a single nanopore run. Mol Ecol Resour 2025;25(1):e14028.

Ivanova NV, Dewaard JR, Hebert PDN. An inexpensive, automation-friendly protocol for recovering high-quality DNA. Mol Ecol Notes 2006;6(4):998–1002.

Kolter A, Gemeinholzer B. Internal transcribed spacer primer evaluation for vascular plant metabarcoding. Metabarcoding Metagenom 2021;5. 10.3897/mbmg.5.68155.

Kress WJ, Erickson DL. A two-locus global DNA barcode for land plants: the coding rbcL gene complements the non-coding trnH-psbA spacer region. PLoS One 2007;2:e508.

Li R, Ratnasingham S, Zarubiieva I et al. PROTAX-GPU: a scalable probabilistic taxonomic classification system for DNA barcodes. Philos Trans R Soc Lond B Biol Sci 2024;379(1904):20230124.

Liu-Wei W, Toorn W van der, Bohn P et al. Sequencing accuracy and systematic errors of nanopore direct RNA sequencing. BMC Genomics 2024;25(1). 10.1186/s12864-024-10440-w.

Lopez JV, Yuhki N, Masuda R et al. Numt, a recent transfer and tandem amplification of mitochondrial DNA to the nuclear genome of the domestic cat. J Mol Evol 1994;39(2):174–90.

Martin M. Cutadapt removes adapter sequences from high-throughput sequencing reads. EMBnet J 2011;17(1):10.

Meier R, Wong W, Srivathsan A et al. $1 DNA barcodes for reconstructing complex phenomes and finding rare species in specimen-rich samples. Cladistics 2016;32(1):100–10.

Meyerhans A, Vartanian JP, Wain-Hobson S. DNA recombination during PCR. Nucleic Acids Res 1990;18(7):1687–91.

Ooms J. Writexl: Export Data Frames to Excel “xlsx” Format. 2025. 10.32614/CRAN.package.writexl.

Pagès H, Aboyoun P, Gentleman R et al. Biostrings: Efficient manipulation of biological strings. Preprint, 2025. 10.18129/B9.bioc.Biostrings.

Pakrashi A, Thompson KA, Hebert PDN. Haplodiploidy accelerates mitogenome evolution in insects. Proc Biol Sci 2025;292(2059):20251813.

Porter TM, Hajibabaei M. MetaWorks: A flexible, scalable bioinformatic pipeline for high-throughput multi-marker biodiversity assessments. PLoS One 2022;17(9):e0274260.

Prosser SWJ, Bard NW, Thompson KA et al. The Metabarcoding Analysis Pipeline (MAP): Simple, accurate, and flexible metabarcoding. In: bioRxiv. bioRxiv, 23 July 2026. 10.64898/2026.07.22.740107.

Prosser SWJ, Floyd RM, Thompson KA et al. BOLDistilled: Automated construction of comprehensive but compact DNA barcode reference libraries. Mol Ecol Resour 2025;25(8):e70043.

Ratnasingham S, Hebert PDN. A DNA-based registry for all animal species: the Barcode Index Number (BIN) system. PLoS One 2013;8(7):e66213.

Rognes T, Flouri T, Nichols B et al. VSEARCH: a versatile open source tool for metagenomics. PeerJ 2016;4(e2584):e2584.

Sahlin K, Lim MCW, Prost S. NGSpeciesID: DNA barcode and amplicon consensus generation from long-read sequencing data. Ecol Evol 2021;11(3):1392–8.

Sánchez-Vendizú P, Erkenswick G, Reyes J et al. Decoding the Peruvian Amazon with in situ DNA barcoding of vertebrate and plant taxa. Sci Data 2025;12(1):1545.

Schnell IB, Bohmann K, Gilbert MTP. Tag jumps illuminated--reducing sequence-to-sample misidentifications in metabarcoding studies. Mol Ecol Resour 2015;15(6):1289–303.

Shagin DA, Shagina IA, Zaretsky AR et al. A high-throughput assay for quantitative measurement of PCR errors. Sci Rep 2017;7(1):2718.

Srivathsan A, Feng V, Suárez D et al. ONTbarcoder 2.0: rapid species discovery and identification with real-time barcoding facilitated by Oxford Nanopore R10.4. Cladistics 2024;40(2):192–203.

Srivathsan A, Lee L, Katoh K et al. ONTbarcoder and MinION barcodes aid biodiversity discovery and identification by everyone, for everyone. BMC Biol 2021;19(1):217.

Tate JA, Simpson BB. Paraphyly of Tarasa (Malvaceae) and diverse origins of the polyploid species. Systematic Botany 2003;28(4):723–37.

Teeling H, Glöckner FO. Current opportunities and challenges in microbial metagenome analysis--a bioinformatic perspective. Brief Bioinform 2012;13(6):728–42.

Wick RR, Judd LM, Holt KE. Deepbinner: Demultiplexing barcoded Oxford Nanopore reads with deep convolutional neural networks. PLoS Comput Biol 2018;14(11):e1006583.

Wickham H, Averick M, Bryan J et al. Welcome to the Tidyverse. Journal of Open Source Software 2019;4(43):1686.

Wickham H, Bryan J. readxl: Read excel files. R Package Version 2019;1(1):785.

